# Fingerprinting of hatchery haplotypes by whole-mitogenome sequencing improves genetic studies of masu salmon *Oncorhynchus masou masou*

**DOI:** 10.1101/2020.01.30.926691

**Authors:** Yoko Kato-Unoki, Keitaro Umemura, Kosuke Tashiro

**Affiliations:** Center for Advanced Instrumental and Educational Supports, Faculty of Agriculture, Kyushu University, Fukuoka, Japan; Fishery Research Laboratory, Kyushu University, Fukuoka, Japan; Laboratory of Molecular Gene Technology, Faculty of Agriculture, Kyushu University, Fukuoka, Japan

**Keywords:** mitochondrial DNA, whole sequence, hatchery fish, indigenous, *Oncorhynchus masou masou*

## Abstract

Stocking hatchery fish can lead to disturbance and extinction of the local indigenous population. Stocking similar lineages to the indigenous population makes it difficult to conduct genetic studies to determine conservation units. We considered that, due to the length of the mitogenome, whole-mitogenome analysis might overcome this problem by enabling identification of hatchery haplotypes. Here, to provide basic information for conservation of indigenous masu salmon *Oncorhynchus masou masou*, a commonly stocked fish in the Kase River system, Japan, we used whole-mitogenome analysis of fish to identify hatchery haplotypes in the river and in several hatcheries that might be used for stocking. Whole-mitogenome sequencing clearly identified hatchery haplotypes like fingerprints, identifying the tributaries contaminated with hatchery haplotypes. The results suggest that informal stocking of *O. m. masou* has been performed widely across the Kase River system. Non-hatchery haplotypes mainly belonged to clade I, which was not found in northern Hokkaido Island. Sites without hatchery haplotypes were estimated three, suggesting that these sites are suitable for conservation of the indigenous fish. This study demonstrated that the resolution of the conventional analysis using partial mitogenome is insufficient to distinguish hatchery haplotype from similar lineages. The whole-mitogenome sequences provided accurate information that was not available in the partial sequences, enabling various inferences: e.g., estimation of the origin of stocked fish and circulation of the reared strains. We conclude that whole-mitogenome analysis is useful for the genetic study of this species. The whole-mitogenome data produced will contribute to the conservation, resource management, and further study of *O. m. masou*.

## Introduction

Stocking of hatchery fish has been carried out throughout the world to mitigate declines in natural production and enhance fishery production and recreational fishing [1]. However, it has become problematic because it can lead to disturbance and extinction of the local indigenous population, and loss of genetic diversity [2,3]. Such effects are observed among salmonid fishes (e.g., [4–7]). Determination of the conservation unit of the indigenous population requires genetic information to identify the indigenous individuals; however, stocking fish with a similar lineage to that of the indigenous fish makes such studies difficult.

Masu salmon *Oncorhynchus masou masou* is endemic across Japan, with diverse life histories [8]: there is a river-resident form and an anadromous form [8], with spawning adults homing to their natal river at high rates, similar to other Pacific salmons [9]. *Oncorhynchus masou masou* is one of commonly stocked fish for recreational fishing in Japan. Especially on Kyushu Island, public resource management is not sufficient in many rivers, and inappropriate and informal stocking is performed widely by individuals and/or fishing clubs. Recent research has suggested that *O. m. masou* can be divided into different populations in terms of behavior, morphology, and genetics in each river [10–13]. Stocking hatchery *O. m. masou* might reduce indigenous characteristics and local adaptations because of hybridization between the stocked and indigenous fish. This could ultimately lead to a loss of indigenous populations, as observed in other salmonid species (e.g., [4,14,15]). To conserve the indigenous *O. m. masou* resource in a river, their genetic information is required. Genetic study of *O. m. masou* in Japan has focused mainly on northern Hokkaido Island and some areas of Honshu Island and its surrounding waters (e.g., [10,13,16,17]); studies of this species in Kyushu Island are scarce [18,19].

The Kase River system is located in northwestern Kyushu Island (Fig 1). Inappropriate *O. m. masou* stocking, with no consideration of the indigenous population, has been conducted for a long time by fishing clubs and personal game fishers in this river. The distribution of this informal stocking and its impacts on indigenous resources are unknown, and genetic information is required.

**Fig 1.**
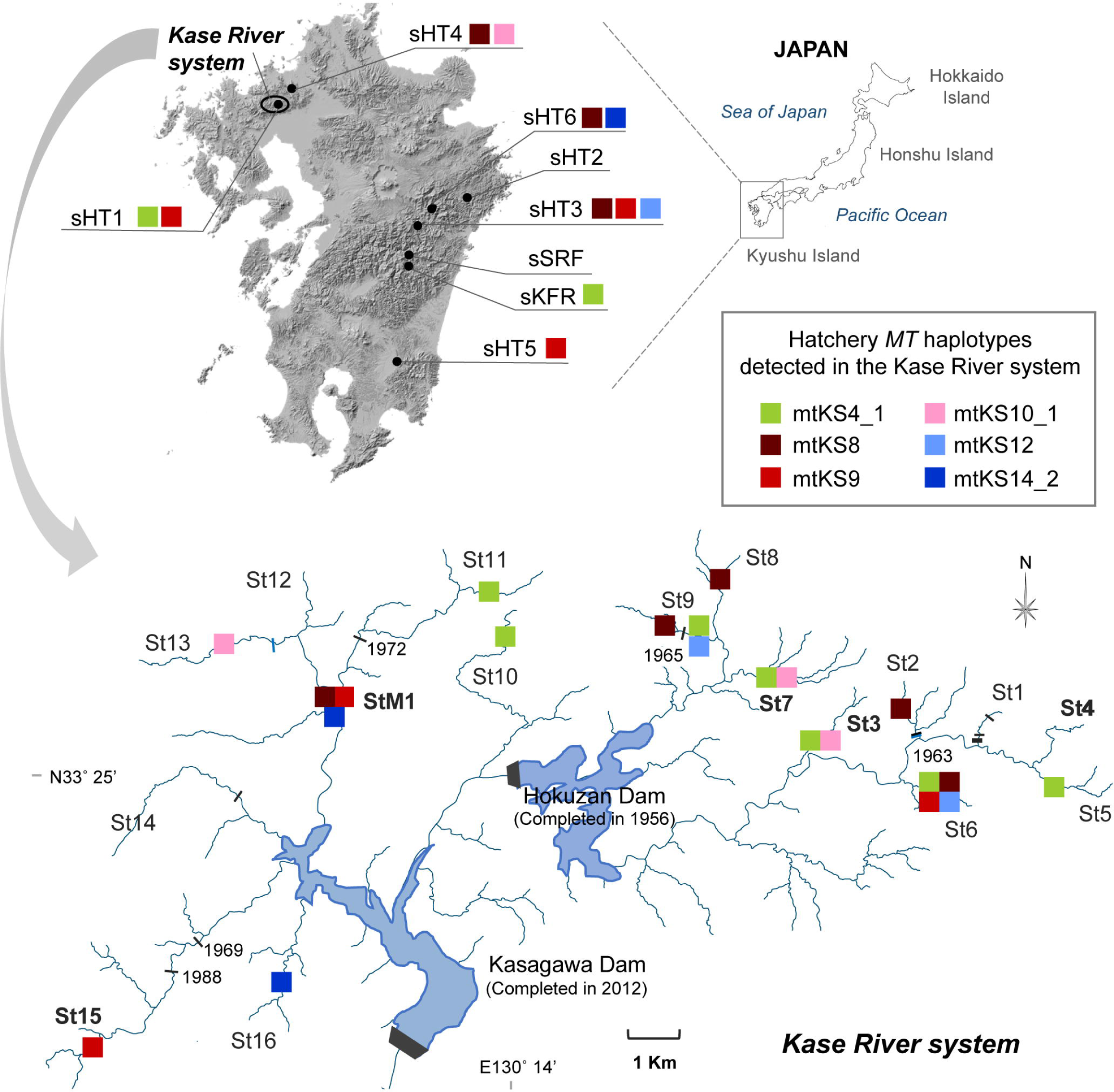
Sampling locations and hatchery *MT* haplotype distribution detected in the Kase River system. The upper map shows Kyushu Island, and the lower map shows the Kase River system. In the lower map, the main man-made dams (black bars with build year) and waterfalls (blue bars) that may hinder the fish run are marked. Sites with prior information about stocking history are show in bold letters. For details of the samples and the results of whole-mitogenome sequencing, see Tables 1 and 4.

We considered that whole-mitogenome analysis might be helpful for genetic studies to determine the conservation units of this indigenous fish. The mitogenome is commonly used as a genetic marker for characterizing population structure and identifying maternal lineages [20–22]. Recent reports suggest that whole-mitogenome analysis can increase the resolution of matrilineal genetic structure patterns, especially in low diversity species, and allow detailed and more accurate phylogenetic analysis (e.g., [23–26]). Thus, whole-mitogenome analysis might enable identification of hatchery haplotype in populations of stocked hatchery fish that share similar lineages, and could help estimate the indigenous genetic structure and lineage. Moreover, whole-mitogenome sequencing will allow comparative phylogenetic analysis of any region of the mitogenome.

**Table 1.** Sample information used in this study.

|  | **Site**<br>**ID** | **River / tributary<sup>a</sup>** | **Sampling**<br>**year** | **Number**<br>**of**<br>**samples** | **Stocking**<br>**history** | **Additional information** |
| --- | --- | --- | --- | --- | --- | --- |
| **Kase River**<br>**system** | St1 | Kase / tributary 1 | 2016 | 16 | unknown |  |
|  | St2 | Kase / tributary 2 | 2016 | 21 | unknown |  |
|  | St3 | Kase / tributary 3 | 2016 | 5 | stocked |  |
|  | St4 | Kase / tributary 4 | 2015, 2016 | 23 | stocked |  |
|  | St5 | Kase / tributary 5 | 2016 | 21 | unknown |  |
|  | St6 | Kase / tributary 6 | 2016 | 12 | unknown |  |
|  | St7 | Kase / tributary 7 | 2016 | 8 | stocked |  |
|  | St8 | Kase / tributary 8 | 2016 | 6 | unknown |  |
|  | St9-1 | Kase / tributary 9 | 2016 | 22 | unknown |  |
|  | St9-2 | Kase / tributary 9 | 2016 | 11 | unknown |  |
|  | St10 | Kase / tributary 10 | 2016 | 26 | unknown |  |
|  | St11 | Kase / tributary 11 | 2016 | 22 | unknown |  |
|  | St12 | Kase / tributary 12 | 2016 | 20 | unknown |  |
|  | St13 | Kase / tributary 13 | 2016 | 16 | unknown |  |
|  | St14 | Kase / tributary 14 | 2016 | 11 | unknown |  |
|  | St15 | Kase / tributary 15 | 2016 | 11 | stocked |  |
|  | St16 | Kase / tributary 16 | 2016 | 11 | unknown |  |
|  | StM1 | Kase / tributary 11 | 2016 | 24 | stocked |  |
| **Control** | sSRF | Hitotsuse | 2016 | 20 | unstocked | prohibited fishing area |
| **Hatchery**<br>**etc.** | sKFR | Hitotsuse / Ishido | 2016 | 7 | - | from native fish [19] |
|  | sHT1 | Hitotsuse / Ishido | 2016 | 11 | - | from sKFR |
|  | sHT2 | Kita | 2016 | 15 | - | from native fish [27] |
|  | sHT3 | Gokase | 2016 | 15 | - | mated with native fish and others<br>[19] |
|  | sHT4 | - | 2016 | 10 | - | mated with many strain |
|  | sHT5 | Ooyodo / Okimizu | 2016 | 10 | - | from native fish [19] |
|  | sHT6 | Gokase / Kawabashiri | 2016 | 9 | - | mated with native fish and others |

<sup>a</sup>For each hatchery, the original river of the reared fish is given.

Here, to provide basic information for conservation of the indigenous *O. m. masou* in the Kase River system, we analyzed the whole-mitogenome in this species in this river system and in several hatcheries that might be used for stocking. Through this analysis we could identify hatchery haplotypes of fish in this river system and clarify their distribution. The whole-mitogenome analysis provided important information that was not available from conventional partial-mitogenome analysis.

## Materials and methods

### Ethics statement

This study was conducted with the permission of the Saga prefecture (permission number 3018). The fish were collected under appropriate fishing licenses that allowed the capture and sacrifice of the fish. The ethical approvals of the Kyushu University Animal Experiment Committee and the Faculty of Agriculture Ethics Committee in the Kyushu University were not required because approval of animal experiment is only necessary for fish reared at Kyushu University (according to the Kyushu University Animal Experimentation Regulations) and ethics covers only human research (this study is fish research) (according to the Faculty of Agriculture Ethics Committee Regulations in the Kyushu University). However, this study was carried out according to the guidelines of the Ichthyological Society of Japan (http://www.fish-isj.jp/english/guidelines.html).

### Study area and samples

Information about the *O. m. masou* samples (sample numbers, sampling year, and stocking history) used in this study is listed in Table 1, and the locations of sampling sites are shown in Fig 1. The samples from the Kase River system were collected at 18 sites from 16 tributaries (St1–St8, St9-1 and St9-2, St10–St16, and StM1) (Fig 1, Table 1). Samples from Shiiba Research Forest, Kyushu University (sSRF), an unstocked site in the Hitotsuse River system, were used as the control group. Information on the stocking history and hatcheries used for the Kase River system were collected, and one research center (Miyazaki Prefectural Fisheries Research Institute, sKFR) and six hatcheries (sHT1–sHT6) were selected as putative sources of *O. m. masou* in this river system (Fig 1, Table 1) [19,27]. Information about the river of origin and mating history of hatchery fishes is shown in the columns “River / tributary” and “Additional information”, respectively, in Table 1.

### Sampling and DNA extraction

Wild fish were captured by electrofishing or normal fishing from 2015 to 2016. A small piece of fin was taken from each specimen, and the captured fish were released at the same point in the river. Tissues were immediately fixed in 99.5% ethanol or RNA*later* solution (Thermo Fisher Scientific) and stored at −20 °C until use. Total genomic DNA was extracted from preserved tissue using the QIAGEN Blood & Tissue DNeasy Kit (QIAGEN).

### DNA sequence and genetic analysis

Sequencing was performed in two steps: first, the NADH dehydrogenase subunit 5 gene (*ND5*) sequences of all samples (383 individuals) were analyzed; next, some of the samples (88 individuals) were selected to represent each site and each *ND5* haplotype, and the whole-mitogenome sequences of these samples were analyzed.

The *ND5* region (1,597 bp) was amplified with the primer pair ND5-F1 and ND5-R (Table 2). The PCR mixture was as follows: 10–50 ng genomic DNA, 1× Phusion HF buffer (New England BioLabs), 0.2 mM of each dNTP, 0.5 µM of each primer, and 0.1 µL Phusion DNA polymerase (New England BioLabs) in a total volume of 10 µL. The PCR program was as follows: one cycle at 98 °C for 30 s, followed by 40 cycles of 98 °C for 10 s, 64 °C for 20 s, and 72 °C for 60 s, and a final extension at 72 °C for 2 min. Each PCR product was sequenced with the primer ND5-F1 or ND5-F2 (Table 2). The *ND5* sequences were assembled and multiple sequences were aligned using ATGC Ver. 4.3.5 software (GENETYX Co.). The identified 1,449 bp sequence variants (*ND5* haplotypes) (*ND5* position, 358–1806; mitogenome position, 13299–14747 for reference sequence NC_008747) were deposited in DDBJ/EMBL/GenBank as shown in S1 Table.

**Table 2.**
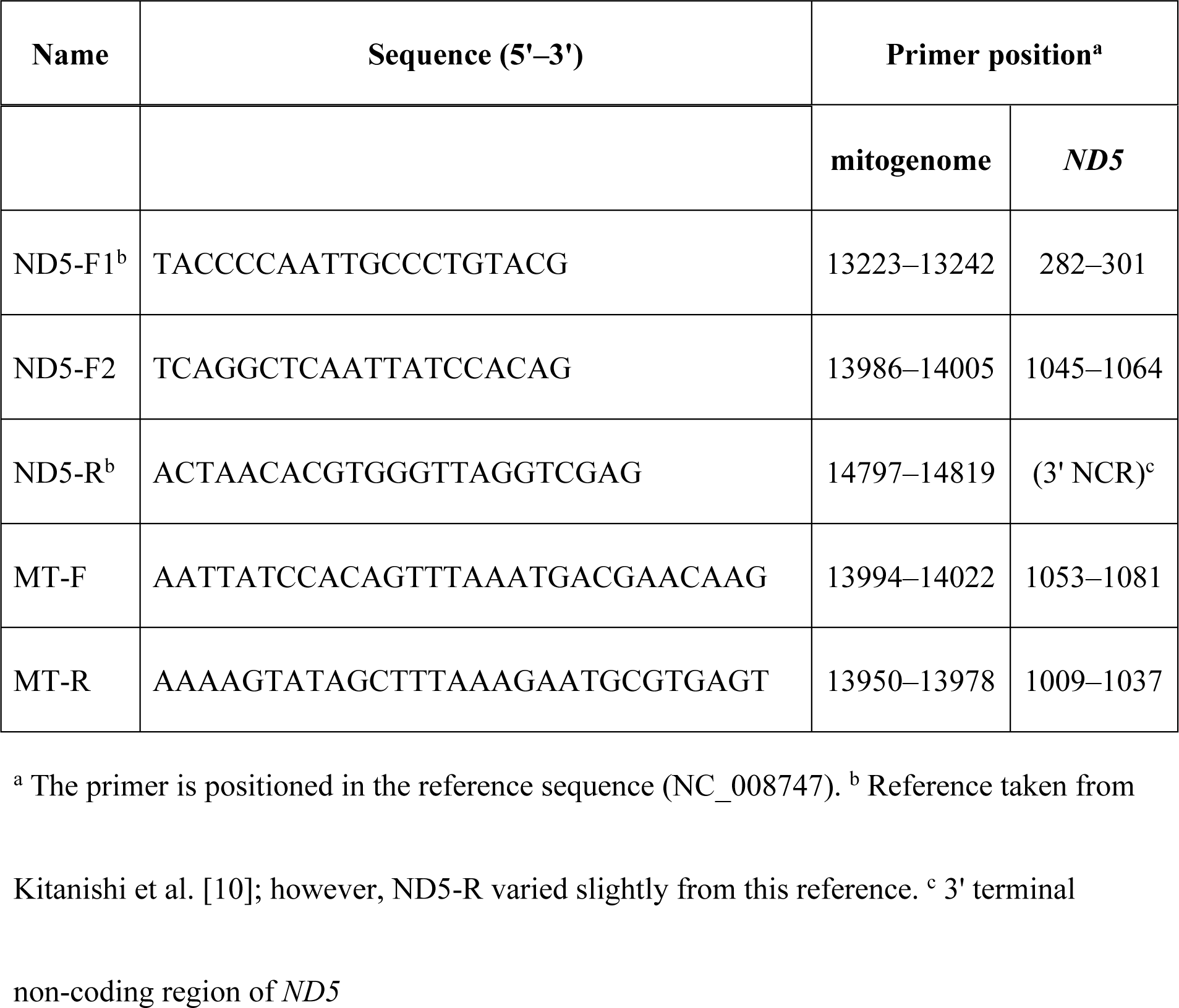
List of primers used in this study.

The whole-mitogenome fragment (16.6 kbp) was amplified with primer pair MT-F and MT-R (Table 2). The PCR mixture was as follows: 20–100 ng genomic DNA, 1× PrimeSTAR GXL Buffer (TaKaRa Bio Inc.), 0.2 mM each dNTP, 0.2 µM each primer, and 0.3 µL of PrimeSTAR GXL DNA polymerase (TaKaRa Bio Inc.) in a total volume of 15 µL. The PCR program was as follows: one cycle at 98 °C for 10 s, followed by 40 cycles of 98 °C for 10 s, 60 °C for 15 s, 68 °C for 14 min, and a final extension at 68 °C for 5 min. Each PCR product was subjected to agarose gel electrophoresis, and the mitogenome was extracted from the excised band by using the Wizard SV Gel and PCR Clean-up System (Promega).

Mitogenome libraries for next-generation sequencing (NGS) were prepared using a QIAseq FX DNA Library Kit (QIAGEN). For each library, the quality and fragment size were checked using an Agilent BioAnalyzer and the concentration was quantified by qPCR (Mx3000p, Agilent) using a KAPA LQ Kit (KAPA Biosystems). Pooled and denatured library (8 pM) containing 5% volume of PhiX (control library; Illumina) was sequenced using the Illumina MiSeq system with MiSeq Reagent Kit V3 (300-bp paired-end reads) (Illumina).

The obtained NGS data of paired end reads were trimmed using Trimmomatic ver. 0.36 as follows: a cleanup adapter was applied and reads with low quality (Q score, <28) and short-length (<50) were filtered out; then, the reads were mapped to the reference mitogenome sequence (*O. m. masou* accession No: NC_008747) using Burrow–Wheeler Aligner ver. 0.7.12, and the obtained SAM (Sequence Alignment Map) files being converted to BAM (Binary Alignment Map) files using SAMtools ver. 1.4.1. The resulting reads and the single nucleotide polymorphism (SNP) sites for the reference sequence were visualized with Integrative Genomics Viewer version 2.3.83. and TASSEL ver. 5.0. The primer sites and those outside the region of interest (mitogenome position: 13950–14022) were replaced with the predetermined *ND5* sequence. The identified full-length mitogenome sequence variants (*MT* haplotypes) were deposited in DDBJ/EMBL/GenBank as shown in S1 Table.

The relationships among haplotypes were estimated using the TCS program implemented in PopART [28]. To determine the relationships between haplotypes in past studies and the current study, we also analyzed the haplotype network including data of Kitanishi et al. [10] and Yu et al. [16].

## Results

### DNA sequence and determination of haplotype

The sequencing analysis of the 1,449 bp fragment containing the ND5 gene detected a total of 18 *ND5* haplotypes with 28 SNP sites from 383 individuals (Tables 3 and S2). Of these, 10 *ND5* haplotypes (KS4, KS6, KS8–KS10, KS12, KS14, HT3, HT4A, and HT4B) were detected in hatcheries, and seven of these, KS4, KS6, KS8–KS10, KS12, and KS14, were detected in the Kase River system. KS4 was also detected in the unstocked area, sSRF, in the Hitotsuse River system.

**Table 3.**
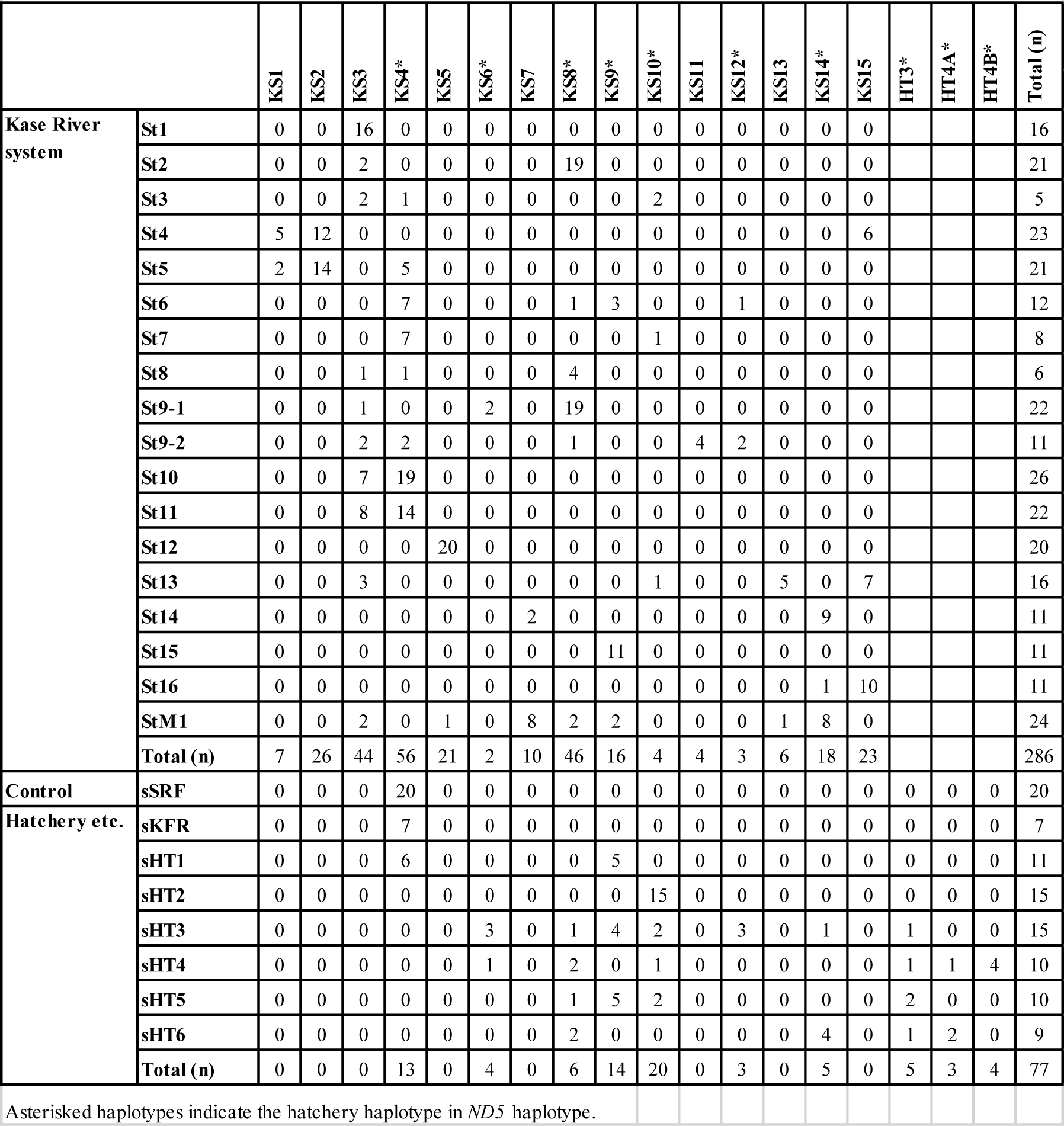
Distribution of *ND5* haplotypes in each sampling site.

|  |  | KS1 | KS2 | KS3 | KS4* | KS5 | KS6* | KS7 | KS8* | KS9* | KS10* | KS11 | KS12* | KS13 | KS14* | KS15 | HT3* | HT4A* | HT4B* | Total (n) |
| --- | --- | --- | --- | --- | --- | --- | --- | --- | --- | --- | --- | --- | --- | --- | --- | --- | --- | --- | --- | --- |
| Kase River system | St1 | 0 | 0 | 16 | 0 | 0 | 0 | 0 | 0 | 0 | 0 | 0 | 0 | 0 | 0 | 0 |  |  |  | 16 |
|  | St2 | 0 | 0 | 2 | 0 | 0 | 0 | 0 | 19 | 0 | 0 | 0 | 0 | 0 | 0 | 0 |  |  |  | 21 |
|  | St3 | 0 | 0 | 2 | 1 | 0 | 0 | 0 | 0 | 0 | 2 | 0 | 0 | 0 | 0 | 0 |  |  |  | 5 |
|  | St4 | 5 | 12 | 0 | 0 | 0 | 0 | 0 | 0 | 0 | 0 | 0 | 0 | 0 | 0 | 6 |  |  |  | 23 |
|  | St5 | 2 | 14 | 0 | 5 | 0 | 0 | 0 | 0 | 0 | 0 | 0 | 0 | 0 | 0 | 0 |  |  |  | 21 |
|  | St6 | 0 | 0 | 0 | 7 | 0 | 0 | 0 | 1 | 3 | 0 | 0 | 1 | 0 | 0 | 0 |  |  |  | 12 |
|  | St7 | 0 | 0 | 0 | 7 | 0 | 0 | 0 | 0 | 0 | 1 | 0 | 0 | 0 | 0 | 0 |  |  |  | 8 |
|  | St8 | 0 | 0 | 1 | 1 | 0 | 0 | 0 | 4 | 0 | 0 | 0 | 0 | 0 | 0 | 0 |  |  |  | 6 |
|  | St9-1 | 0 | 0 | 1 | 0 | 0 | 2 | 0 | 19 | 0 | 0 | 0 | 0 | 0 | 0 | 0 |  |  |  | 22 |
|  | St9-2 | 0 | 0 | 2 | 2 | 0 | 0 | 0 | 1 | 0 | 0 | 4 | 2 | 0 | 0 | 0 |  |  |  | 11 |
|  | St10 | 0 | 0 | 7 | 19 | 0 | 0 | 0 | 0 | 0 | 0 | 0 | 0 | 0 | 0 | 0 |  |  |  | 26 |
|  | St11 | 0 | 0 | 8 | 14 | 0 | 0 | 0 | 0 | 0 | 0 | 0 | 0 | 0 | 0 | 0 |  |  |  | 22 |
|  | St12 | 0 | 0 | 0 | 0 | 20 | 0 | 0 | 0 | 0 | 0 | 0 | 0 | 0 | 0 | 0 |  |  |  | 20 |
|  | St13 | 0 | 0 | 3 | 0 | 0 | 0 | 0 | 0 | 0 | 1 | 0 | 0 | 5 | 0 | 7 |  |  |  | 16 |
|  | St14 | 0 | 0 | 0 | 0 | 0 | 0 | 2 | 0 | 0 | 0 | 0 | 0 | 0 | 9 | 0 |  |  |  | 11 |
|  | St15 | 0 | 0 | 0 | 0 | 0 | 0 | 0 | 0 | 11 | 0 | 0 | 0 | 0 | 0 | 0 |  |  |  | 11 |
|  | St16 | 0 | 0 | 0 | 0 | 0 | 0 | 0 | 0 | 0 | 0 | 0 | 0 | 0 | 1 | 10 |  |  |  | 11 |
|  | StM1 | 0 | 0 | 2 | 0 | 1 | 0 | 8 | 2 | 2 | 0 | 0 | 0 | 1 | 8 | 0 |  |  |  | 24 |
|  | Total (n) | 7 | 26 | 44 | 56 | 21 | 2 | 10 | 46 | 16 | 4 | 4 | 3 | 6 | 18 | 23 |  |  |  | 286 |
| Control | sSRF | 0 | 0 | 0 | 20 | 0 | 0 | 0 | 0 | 0 | 0 | 0 | 0 | 0 | 0 | 0 | 0 | 0 | 0 | 20 |
| Hatchery etc. | sKFR | 0 | 0 | 0 | 7 | 0 | 0 | 0 | 0 | 0 | 0 | 0 | 0 | 0 | 0 | 0 | 0 | 0 | 0 | 7 |
|  | sHT1 | 0 | 0 | 0 | 6 | 0 | 0 | 0 | 0 | 5 | 0 | 0 | 0 | 0 | 0 | 0 | 0 | 0 | 0 | 11 |
|  | sHT2 | 0 | 0 | 0 | 0 | 0 | 0 | 0 | 0 | 0 | 15 | 0 | 0 | 0 | 0 | 0 | 0 | 0 | 0 | 15 |
|  | sHT3 | 0 | 0 | 0 | 0 | 0 | 3 | 0 | 1 | 4 | 2 | 0 | 3 | 0 | 1 | 0 | 1 | 0 | 0 | 15 |
|  | sHT4 | 0 | 0 | 0 | 0 | 0 | 1 | 0 | 2 | 0 | 1 | 0 | 0 | 0 | 0 | 0 | 1 | 1 | 4 | 10 |
|  | sHT5 | 0 | 0 | 0 | 0 | 0 | 0 | 0 | 1 | 5 | 2 | 0 | 0 | 0 | 0 | 0 | 2 | 0 | 0 | 10 |
|  | sHT6 | 0 | 0 | 0 | 0 | 0 | 0 | 0 | 2 | 0 | 0 | 0 | 0 | 0 | 4 | 0 | 1 | 2 | 0 | 9 |
|  | Total (n) | 0 | 0 | 0 | 13 | 0 | 4 | 0 | 6 | 14 | 20 | 0 | 3 | 0 | 5 | 0 | 5 | 3 | 4 | 77 |

Asterisked haplotypes indicate the hatchery haplotype in *ND5* haplotype.

**Table 4.**
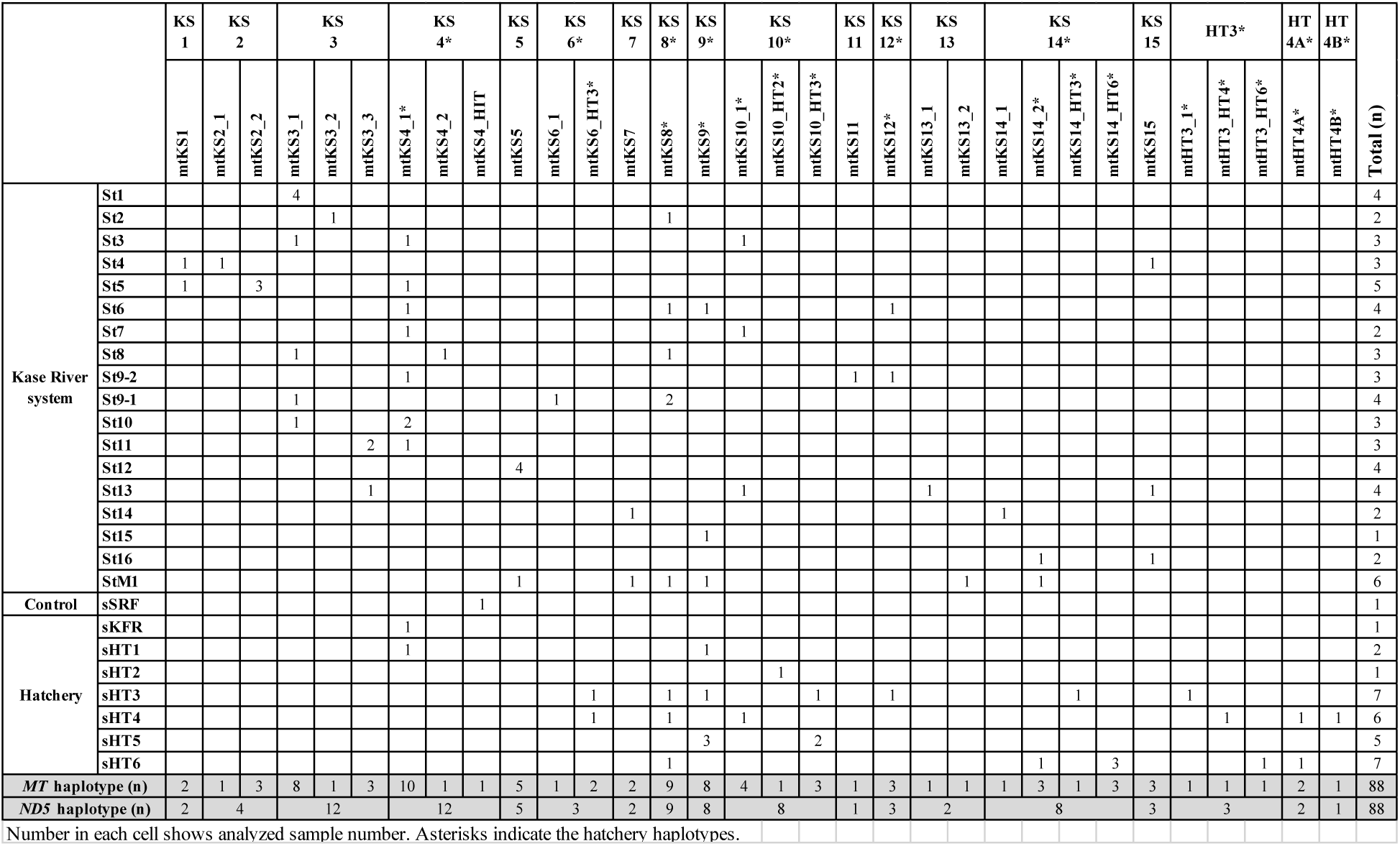
Result of whole-mitogenome sequencing.

Eighty-eight of the 383 individuals that underwent *ND5* sequencing were selected for whole-genome sequencing by NGS. The whole-mitogenome sequencing of all 88 individuals was successful with approximately >1500-fold depth of coverage. The number of samples with each determined *MT* haplotype are shown for each sampling site in Table 4. The majority of *ND5* haplotypes were associated with more than one *MT* haplotype, with 32 haplotypes including 256 SNP sites being detected (Tables 4 and S2). Sixteen haplotypes were detected in hatcheries (see asterisks, Table 4), and six of these, mtKS4_1, mtKS8, mtKS9, mtKS10_1, mtKS12, and mtKS14_2, were detected in the Kase River system. Hatchery haplotypes were distributed in 12 tributaries (13 sites) including sites with unknown stocking history (Fig 1). Several hatchery haplotypes were detected upstream of artificial barriers (e.g., dams) and/or natural barriers (e.g., waterfalls): mtKS4_1 at St11; mtKS8 at St2 and St9-1; mtKS9 at St15; and mtKS10_1 at St13 (Fig 1). The whole-mitogenome sequences of KS6 in the Kase River system (mtKS6_1) and KS4 in sSRF (mtKS4_2) did not match any hatchery *MT* haplotype in the samples analyzed here (Table 4). Two *MT* haplotypes, mtKS4_1 and mtKS9, were detected in hatchery sHT1, which was reared with fish from sKFR; one of these *MT* haplotypes, mtKS4_1, was the same as that detected in sKFR (Tables 1 and 4). Analyzed MT haplotype in the ND5 haplotype KS4 samples in the Kase River system was consistently mtKS4_1 except a sample at St8, mtKS4_2.

Some hatchery haplotypes were detected at multiple hatcheries (Table 4): mtKS6_HT3 was detected at sHT3 and sHT4; mtKS8 was detected at sHT3, sHT4, and sHT6; mtKS9 was detected at sHT1, sHT3, and sHT5; mtKS10_HT3 was detected at sHT3 and sHT5; and mtHT4A was detected at sHT4 and sHT6. On the other hand, seven hatchery haplotypes (mtKS10_HT2, mtKS14_HT3, mtKS14_HT6, mtHT3_1, mtHT3_HT4, mtHT3_HT6, and mtHT4B) were not detected in any of the samples from the Kase River system and other hatcheries.

### Haplotype network analysis

We analyzed the relationships among haplotypes by using the TCS program (see Materials and Methods). The results showed a network of *ND5* haplotypes (*ND5*-haplotype network) comprising four clades (Fig 2A). Analysis of *MT* haplotype relationships showed a detailed network (*MT*-haplotype network) (Fig 2B). Clades I–III in the *ND5*-haplotype network were similar to those in the *MT*-haplotype network. Clade IV of the *ND5*-haplotype network included other *O. masou* subspecies and formed a star-like topology radiating from haplotype KS14, whereas clade IV of the *MT*-haplotype network was divided into several sub-clades. In both the *ND5-* and *MT*-haplotype networks, all haplotypes in clade III were hatchery haplotypes. In clade I of the *MT*-haplotype network, mtKS4_1, which was detected at sKFR (original river: Ishido tributary in the Hitotsuse River system), was in the same branch as mtKS4_HIT, which was detected at sSRF (original river: other tributary in the Hitotsuse River system).

**Fig 2.**
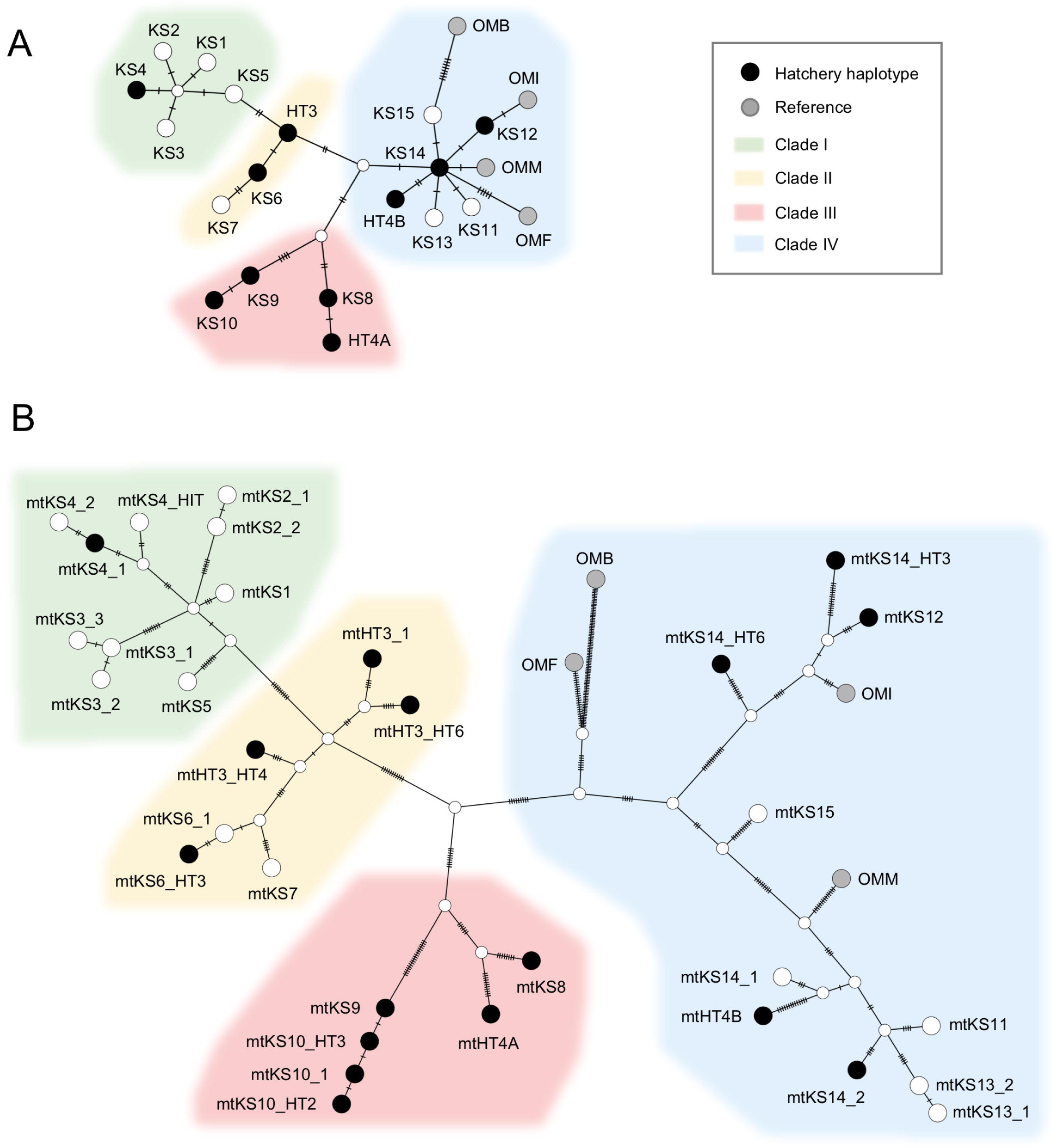
TCS network trees of *ND5* haplotypes and *MT* haplotypes obtained in this study. TCS network trees of (A) *ND5* haplotypes and (B) *MT* haplotypes are shown. Each dash represents one single nucleotide difference between two neighboring haplotypes. Reference information (abbreviations and sequence accession numbers) are as follows: OMM, *O. m. masou* (NC_008747); OMI, *O. m. ishikawae* (DQ_864464); OMF, *O. m. formosanus* (DQ_858456); OMB, *O. m*. Biwa subsp. (Biwa salmon) (EF_105342).

In the *ND5*-haplotypes network, the non-hatchery haplotypes were mostly in clade I (70%), followed by clade IV (23%) and clade II (7%) (from the data in Table 3). We compared the haplotypes and cladal relationships obtained in this study with those obtained using sequencing data of a 561 bp region in *ND5* in the reports by Kitanishi et al. [10] and Yu et al. [16] (S3 Table and S1 Fig). KS1, KS2, and KS13, were new haplotypes in this region of *ND5* in the current study. Clades I, II, III, and IV in the current study appear to correspond to clade 1-3, 1-6, 2-3, and 2-1, reported by Yu et al. [16], respectively.

## Discussion

### Identification of hatchery haplotypes by whole-mitogenome analysis

The whole-mitogenome sequencing clearly identified hatchery haplotypes in *O. m. masou*, like fingerprints. Tributaries contaminated with fish with hatchery haplotypes were clarified, suggesting that informal stocking was widely done across the study area. In pre-information, five tributaries (five sites) had a stocking history (Table 1; bold letters, Fig 1), however, hatchery haplotypes were detected in 12 tributaries (13 sites) (Fig 1). Hatchery haplotypes were also detected upstream of waterfalls and artificial barriers that were built before stocking become popular in the Kase River system in the early 1970s (Fig 1)[29]. *Oncorhynchus masou masou* has a high homing tendency for the natal stream, and the dispersal of anadromous females and river-resident individuals is low [30–32]. Hence, we consider that hatchery haplotypes in each tributary were derived from artificially stocked fish rather than natural dispersal.

This study suggests that the resolution of partial mitogenome sequences is insufficient to distinguish between hatchery haplotypes and indigenous haplotypes in similar lineages. For example, the *ND5* haplotype detected in the prohibited fishing area sSRF was identical to that detected at the hatcheries sHT1 and sKFR (KS4; Table 3), however, the *MT* haplotype detected at sSRF (i.e., mtKS4_HIT) and that detected at the above hatcheries (i.e., mtKS4_1) differed by four nucleotides (Tables 4 and S2). Without whole-mitogenome analysis, we could not get the accurate understanding of the result indicating that the samples in the prohibited fishing area showed a hatchery *ND5* haplotype, as described in next paragraph.

Our results suggest that the *MT* haplotype mtKS4_HIT might be an indigenous haplotype in sSRF for the following reasons. As mentioned above, the mtKS4_HIT haplotype detected in sSRF differed by only four nucleotides from the mtKS4_1 haplotype found in the sHT1 hatchery and sKFR, and they diverged from the same node in the *MT*-haplotype network (Fig 2B). Fish in the sHT1 hatchery are known to originate from sKFR, which in turn sourced fish from the Ishido tributary of the Hitotsuse River system. The site sSRF is in another tributary of the same Hitotsuse River system (Table 1). Therefore it is possible that fish with these two haplotypes diverged from a common ancestor in the Hitotsuse River system and accumulated genetic mutations at each site due to their homing ability and/or being landlocked [32,33]. These results also indicate that the whole-mitogenome analysis can detect a subtle divergence of the mitogenome among tributaries.

### Other findings provided by whole-mitogenome analysis

The whole-mitogenome analysis provided accurate information that was not available by conventional partial analysis of the mitogenome, enabling various inferences to be made. For instance, it allowed estimation of the origin of stocked fish. Partial analysis showed that the *ND5* haplotype KS4 was detected widely and frequently across the Kase River system (Table 3). However, with whole-mitogenome analysis, fish with this haplotype could be divided into three *MT* haplotypes, mtKS4_1, mtKS4_2, and mtKS4_HIT, and these *MT* haplotype of analyzed samples in the Kase River system were all mtKS4_1 except a sample at St8, mtKS4_2 (Table 4). Hence, we could infer that the haplotype mtKS4_1 in the Kase River system was derived from sKFR and was spread from the hatchery sHT1 on the Kase River system, because this *MT* haplotype was detected at both sHT1 and its original sKFR, consistent with our pre-information (Fig 1, Table 1).

The current study clarified that a reared strain is shared by multiple hatcheries and is stocked at multiple sites: e.g., hatchery *MT* haplotype mtKS8 was detected at three hatcheries (sHT3, sHT4, and sHT6) and four tributaries (five sites: St2, St6, St8, St9-1, and StM1) (Fig 1, Table 4). Hatcheries sHT2–sHT6 are used by fisheries cooperatives, fishing clubs, and/or many personal people are likely used to stock various rivers (e.g., [19,27]). It is of concern that stocking of these rivers could cause loss of the unique gene sequences of the indigenous fish, and lead to loss of genetic diversity in this species in the river and/or tributaries [2,14,34].

Non-hatchery haplotypes habitats are estimated to three sites. Urgent conservation might be required. Indigenous fish might be remained in the tributaries in which we did not detect a hatchery haplotype, and no hatchery haplotypes were detected four tributaries (4 sites) in the Kase River system (sites St1, St4, St12 and St14; Fig 1 and Table 4). However, the mtKS15 haplotype detected at St4 could be a stocked hatchery haplotype that could not be detected in this study, because (a) St4 has a stocking record (Table 1), (b) mtKS15 was also detected from the sites, St13 and St16, which are geographically distant from St4, as observed with other hatchery *MT* haplotypes. (Fig 1, Table 4). Indigenous fish in the Kase River system may be revealed in further studies by microsatellite analysis of the nucleic genome [22,35]. Addition, this suspicious haplotype might be resolved by accumulating whole-mitogenome sequence data of hatcheries and field samples from several other areas in Japan in future studies.

### Future applications of whole-mitogenome analysis

The SNP data obtained here for *O. m. masou* (S2 Table) may be used for resource management such as monitoring of hatchery fishes and/or putative indigenous fish by testing environmental DNA [36,37].

The whole-mitogenome analysis will help estimate the indigenous haplotype, because the indigenous haplotypes must be among the non-hatchery haplotypes. Estimation of the indigenous haplotype might lead to research into the geographical and phylogenetic evolution of this species. In this study, the proportion of non-hatchery haplotypes was highest in clade I (70%), possibly suggesting that clade I is the main population of indigenous *O. m. masou* in the Kase River system. As shown in S1 Fig, we obtained three new haplotypes, KS1, KS2, and KS13, in the *ND5* 561 bp region in this study. KS1 and KS2 are not hatchery haplotypes and belong to clade I (Fig 2). The only haplotype corresponding to clade I in previous studies is H22 (S3 Table and S1 Fig) [16], however this haplotype was not detected in northern Hokkaido Island and was detected at a low rate in Honshu Island and in Korea [10,16]. These results may suggest that the clade composition differs between the Kase River system on Kyushu Island and northern Japan. A recent study using a cytochrome-*b* gene (1141 bp) in *O. masou* in the northwestern Pacific including southern Kyushu Island suggested the possible existence of a specific clade comprised of only *O. m. masou* in Kyushu Island (Group D in [18]). Further study might be necessary to assess whether our clade I haplotypes are related to Group D haplotypes in that report. Accumulating the whole-mitogenome data for this species throughout its area of distribution will might allow for a detailed phylogeographic analysis and may reveal the history of geographic evolution in this species.

## Acknowledgments

The authors are grateful to Dr. Norio Onikura of the Faculty of Agriculture of Kyushu University for supporting this study; Dr. Yukiko Ogino of the Faculty of Agriculture of Kyushu University for their help with English writing; Dr. Takamasa Kawamura of the Faculty of Agriculture of Kyushu University for technical support of NGS; Daisuke Kambayashi of Miyazaki Prefectural Fisheries Research Institute and staff at the Shiiba Research Forest of Kyushu University for providing samples; and the Fisheries Division, River & Sediment Control Division, and Forestry Preservation Division of Saga Prefecture for providing information. NGS was performed using an Illumina MiSeq at the Center for Advanced Instrumental and Educational Support, Faculty of Agriculture, Kyushu University. This work was supported by the Japan Society for the Promotion of Science KAKENHI Grant Number 16H00474.

## Author Contributions

Conceptualization: YK.

Formal analysis: YK, KU, KT.

Funding acquisition: YK.

Investigation: YK, KU, KT.

Methodology: YK.

Project administration: YK.

Resources (NGS reagents): KT.

Supervision: YK.

Writing – original draft: YK.

Writing – review & editing: KU, KT.

## Supporting information captions

**S1 Table. List of GenBank accession numbers deposited in this study**.

**S2 Table. List of *MT* haplotype SNP sequences for the reference sequence (*O. m. masou* NC_008747)**.

**S3 Table. Collation of haplotypes determined in past studies and this study**.

**S1 Fig. TCS network trees showing the relationship between data from past studies and the present study**. Double circled haplotypes are new haplotypes obtained by this study. Colors show the clades corresponded to this study in Fig 2. For details showing the collation of haplotypes of past studies and this study see S3 Table.

## References

1. Lever C. Naturalized Fishes of the World. London: Academic Press; 1996. Available: https://books.google.co.jp/books?id=Q5QWAQAAIAAJ

2. Hindar K, Ryman N, Utter F. Genetic Effects of Cultured Fish on Natural Fish Populations. Can J Fish Aquat Sci. 1991;48: 945–957.

3. Rhymer JM, Simberloff D. Extinction by hybridization and introgression. Annu Rev Ecol Syst. 1996;27: 83–109. doi: 10.1146/annurev.ecolsys.27.1.83

4. Araki H, Berejikian BA, Ford MJ, Blouin MS. Fitness of hatchery-reared salmonids in the wild. Evol Appl. 2008;1: 342–355. doi: 10.1111/j.1752-4571.2008.00026.x

5. Altukhov YP, Salmenkova EA. Stock transfer relative to natural organization, management, and conservation of fish populations. In: Ryman N, Utter FM, editors. Population genetics and fishery management. Seattle: University of Washington Press; 1987. pp. 333–343.

6. Miller LM, Close T, Kapuscinski AR. Lower fitness of hatchery and hybrid rainbow trout compared to naturalized populations in Lake Superior tributaries. Mol Ecol. 2004;13: 3379–3388. doi: 10.1111/j.1365-294X.2004.02347.x

7. Gharrett AJ, Smoker WW, Reisenbichler RR, Taylor SG. Outbreeding depression in hybrids between odd- and even-broodyear pink salmon. Aquaculture. 1999;173: 117–129. doi: https://doi.org/10.1016/S0044-8486(98)00480-3

8. Kato F. Life histories of masu and amago salmon (*Oncorhynchus masou* and *Oncorhynchus rhodurus*). In: Groot C, Margolis L, editors. Pacific Salmon Life Histories. Vancouver: University of British Columbia Press; 1991. pp. 396–414. Available: http://ci.nii.ac.jp/naid/10008268078/en/

9. Ueda H. Sensory mechanisms of natal stream imprinting and homing in *Oncorhynchus* spp. J Fish Biol. 2019;95: 293–303. doi: 10.1111/jfb.13775

10. Kitanishi S, Edo K, Yamamoto T, Azuma N, Hasegawa O, Higashi S. Genetic structure of masu salmon (*Oncorhynchus masou*) populations in Hokkaido, northernmost Japan, inferred from mitochondrial DNA variation. J Fish Biol. 2007;71: 437–452. doi: 10.1111/j.1095-8649.2007.01689.x

11. Mayama H. *Oncorhynchus masou masou* ecological notes (in Japanese). Fish Egg. 1990;159: 7–21. Available: http://salmon.fra.affrc.go.jp/kankobutu/tech_repo/fe02/fishandegg159_p07-21.pdf

12. Noguchi D, Taniguch N. Studies on the genetic diversity of wild populations by microsatellite of masu *Oncorhynchus* DNA markers (in Japanese with English summary). Aquac Sci. 2007;55: 521–527. doi: 10.10233/aquaculturesci1953.55.521

13. Kitanishi S, Yamamoto T, Urabe H, Shimoda K. Hierarchical genetic structure of native masu salmon populations in Hokkaido, Japan. Environ Biol Fishes. 2018;101: 699–710. doi: 10.1007/s10641-018-0730-6

14. Finnegan AK, Stevens JR. Assessing the long-term genetic impact of historical stocking events on contemporary populations of Atlantic salmon, *Salmo salar*. Fish Manag Ecol. 2008;15: 315–326. doi: 10.1111/j.1365-2400.2008.00616.x

15. Hansen MM, Fraser DJ, Meier K, Mensberg KLD. Sixty years of anthropogenic pressure: A spatio-temporal genetic analysis of brown trout populations subject to stocking and population declines. Mol Ecol. 2009;18: 2549–2562. doi: 10.1111/j.1365-294X.2009.04198.x

16. Yu J, Azuma N, Yoon M, Brykov V, Urawa S, Nagata M, et al. Population genetic structure and phylogeography of masu salmon (*Oncorhynchus masou masou*) inferred from mitochondrial and microsatellite DNA analyses. Zoolog Sci. 2010;27: 375–385. doi: 10.2108/zsj.27.375

17. Yamamoto S, Morita K, Kikko T, Kawamura K, Sato S, Gwo J-C. Phylogeography of a salmonid fish, masu salmon *Oncorhynchus masou* subspecies-complex, with disjunct distributions across the temperate northern Pacific. Freshw Biol. 2019; 1–18. doi: 10.1111/fwb.13460

18. Iwatsuki Y, Ineno T, Tanaka F, Tanaka K. The southernmost population of *Onchorhynchus masou masou* from Kyushu Island, Japan and gross genetic structure of the *O. masou* complex from the northwestern Pacific. In: Gwo J-C, Shieh Y-T, Burridge CP, editors. Proceedings of International Symposium on the 100th Anniversary of the discovery of Formosa landlocked salmon Bull Natl Taiwan Museum. National Taiwan Museum; 2019. pp. 101–119.

19. Ineno T, Taguchi T. A study on the genetic diversity of inland waters fish and shellfish. Bull MiyazakiPref Fish Exp Stn. 2007;1: 311-317 (in Japanese). Available: http://www.mz-suishi.jp/cgi-bin/upload22/0232_Taro_0404.pdf

20. Cann RL, Stoneking M, Wilson AC. Mitochondrial DNA and human evolution. Nature. 1987;325: 31–36. doi: 10.1038/325031a0

21. Forgacs D, Wallen RL, Dobson LK, Derr JN. Mitochondrial Genome Analysis Reveals Historical Lineages in Yellowstone Bison. PLoS One. 2016;11: e0166081. Available: https://doi.org/10.1371/journal.pone.0166081

22. Fraser DJ, Bernatchez L. Adaptive evolutionary conservation: Towards a unified concept for defining conservation units. Mol Ecol. 2001;10: 2741–2752. doi: 10.1046/j.1365-294X.2001.t01-1-01411.x

23. Miya M, Nishida M. Use of mitogenomic information in teleostean molecular phylogenetics: a tree-based exploration under the maximum-parsimony optimality criterion. Mol Phylogenet Evol. 2000;17: 437–455.

24. Shamblin BM, Bjorndal KA, Bolten AB, Hillis-Starr ZM, Lundgren I, Naro-Maciel E, et al. Mitogenomic sequences better resolve stock structure of southern Greater Caribbean green turtle rookeries. Mol Ecol. 2012;21: 2330–2340. doi: 10.1111/j.1365-294X.2012.05530.x

25. Feutry P, Kyne PM, Pillans RD, Chen X, Marthick JR, Morgan DL, et al. Whole mitogenome sequencing refines population structure of the Critically Endangered sawfish Pristis pristis. Mar Ecol Prog Ser. 2015;533: 237–244. doi: 10.3354/meps11354

26. Bishop CR, Hughes JM, Schmidt DJ. Mitogenomic analysis of the Australian lungfish (*Neoceratodus forsteri*) reveals structuring of indigenous riverine populations and late Pleistocene movement between drainage basins. Conserv Genet. 2017;19: 587–597. doi: 10.1007/s10592-017-1034-7

27. Kimoto K, Mekata T, Takahashi H, Nagasawa K. Genetic structure of the amago and iwame forms of the red-spotted masu salmon *Oncorhynchus masou ishikawae* in the upper Ono River, northeastern Kyushu, southern Japan. Aquacult Sci. 2015;63: 299–309. doi: 10.10233/aquaculturesci.63.299

28. Leigh JW, Bryant D. POPART: Full-feature software for haplotype network construction. Methods Ecol Evol. 2015;6: 1110–1116. doi: 10.1111/2041-210X.12410

29. Kimura S. *Oncorhynchus masou masou* in Kyushu Ialand, specially reference to its spawning behavior. Animals of Kyusyu and Okinawa, 2. Nishinippon Shinbunsha; 1976. pp. 36-63 (in Japanese).

30. Miyakoshi Y, Takahashi M, Ohkuma K, Urabe H, Shimoda K, Kawamuka H. Homing of masu salmon in the tributaries of the Shiribetsu River evaluated by returns of marked fish (in Japanese with English summary). Sci Rep Hokkaido Fish Res Inst. 2012;81: 125–129. Available: https://www.hro.or.jp/list/fisheries/marine/att/81-miyakoshi1.pdf

31. Ueda H. Physiological mechanisms of imprinting and homing migration in Pacific salmon *Oncorhynchus* spp. J Fish Biol. 2012;81: 543–558. doi: 10.1111/j.1095-8649.2012.03354.x

32. Kitanishi S, Yamamoto T, Koizumi I, Dunham JB, Higashi S. Fine scale relationships between sex, life history, and dispersal of masu salmon. Ecol Evol. 2012;2: 920–929. doi: 10.1002/ece3.228

33. Kitanishi S, Yamamoto T, Higashi S. Microsatellite variation reveals fine-scale genetic structure of masu salmon, *Oncorhynchus masou*, within the Atsuta River. Ecol Freshw Fish. 2009;18: 65–71. doi: 10.1111/j.1600-0633.2008.00325.x

34. Lodge DM. Biological Invasions - lessons for ecology. Trends Ecol Evol. 1993;8: 133–137. doi: 10.1016/0169-5347(93)90025-K

35. Toews DPL, Brelsford A. The biogeography of mitochondrial and nuclear discordance in animals. Mol Ecol. 2012;21: 3907–3930. doi: 10.1111/j.1365-294X.2012.05664.x

36. Rees HC, Maddison BC, Middleditch DJ, Patmore JRM, Gough KC. REVIEW The detection of aquatic animal species using environmental DNA - a review of eDNA as a survey tool in ecology. J Appl Ecol. 2014;51: 1450–1459. doi: 10.1111/1365-2664.12306

37. Tsuji S, Maruyama A, Miya M, Ushio M, Sato H, Minamoto T, et al. Environmental DNA analysis shows high potential as a tool for estimating intraspecific genetic diversity in a wild fish population. bioRxiv. 2019; 1–24. doi: http://dx.doi.org/10.1101/829770

